## supplemental figures for "Heterogeneous presynaptic receptive fields contribute to directional tuning in starburst amacrine cells"

**Supplementary**

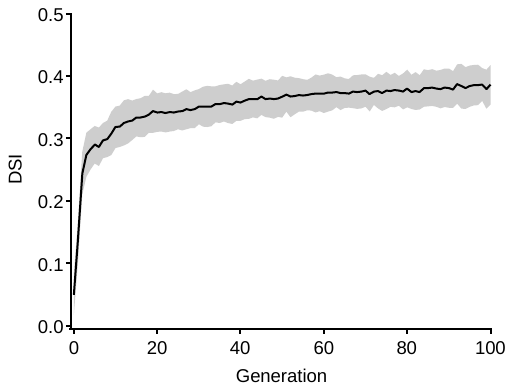

**Figure 2 S1: Example evolution of the directional tuning in a bipolar-SAC model.**

DSI measured from the best performing model obtained through evolutionary algorithm training.

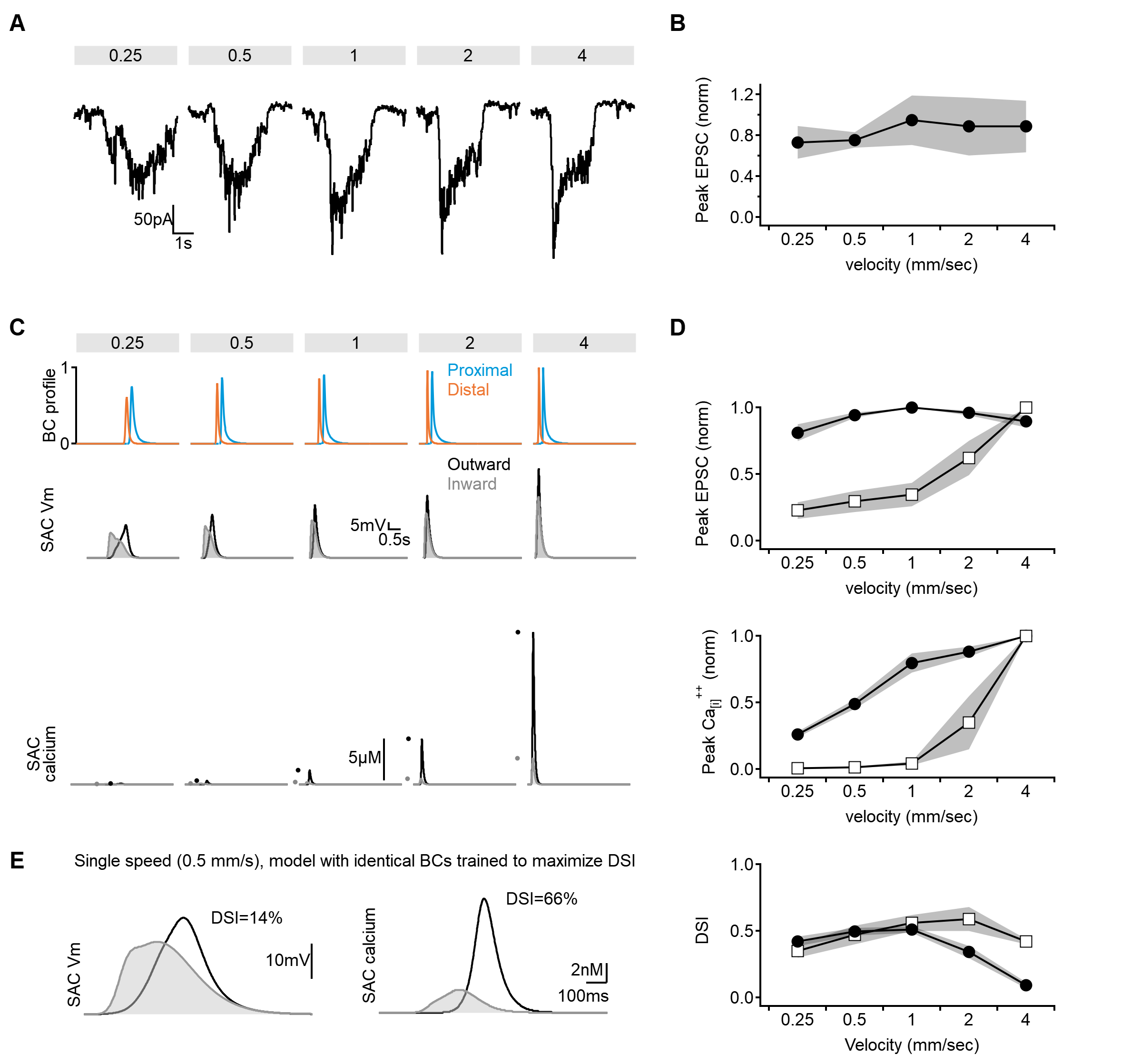

**Figure 2 S2: Limitations of model training on DSI alone.**

**A, B**) Tuning curves of experimentally recorded excitatory inputs in SACs. **A**) representative EPSC profile, obtained by averaging three responses from the same cell). **B**) The normalized peak EPSC amplitudes (shaded areas - SD, n = 4).

**C**) Example profiles of presynaptic bipolar release (top), corresponding voltages (middle), and calcium responses (bottom) in SAC dendrite from models trained to maximize the DSI values.

**D**) Comparison of the peak somatic currents (top), dendritic calcium levels (middle), and DSI (bottom) as a function of stimulus velocity in model formulation presented in **Figure 2** (training objective was large DSI and strong calcium responses, black circles) vs. the model trained on maximizing DSI only (as in **C**, open squares). The model trained solely on DSI fails to capture the observed velocity profiles of the somatic currents and the reported stability of calcium levels.

**E**) Models trained to maximize the DSI of responses to single velocity converge on solutions that depend on string nonlinearities near the activation threshold of the calcium channel. In these conditions, pronounced DS can be observed even with identical BC formulations over the whole SAC dendritic tree.

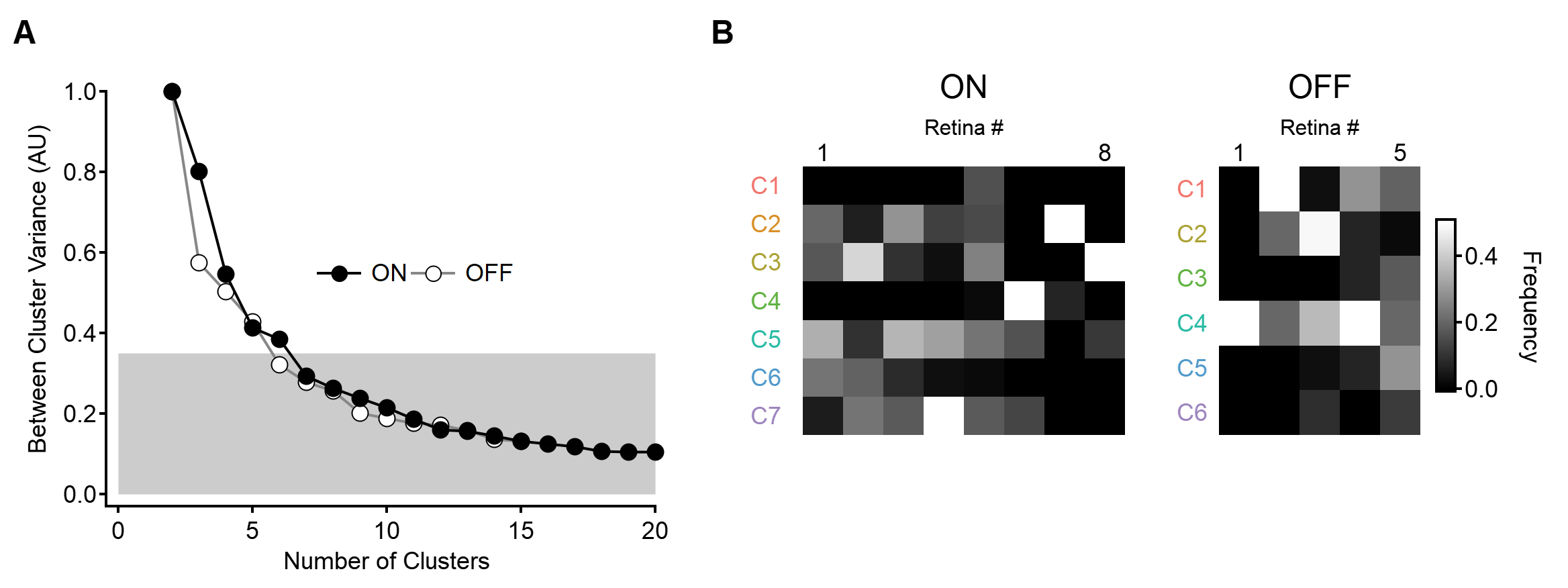

**Figure 6-S1: Determining the optimal number of functional clusters for the ON- and OFF-SAC populations.**

**A**) The variance between input clusters computed based on the shapes of motion responses in individual ROIs, as the function of the number of clusters. The metric was normalized to the levels observed with 2 clusters. The number of clusters was selected as the threshold required to cross e^-1^ (~36%) of unexplained variance.

B) The prevalence of the functional clusters across recordings. Color coding indicates the relative frequency of the functional clusters observed within each retina preparation.

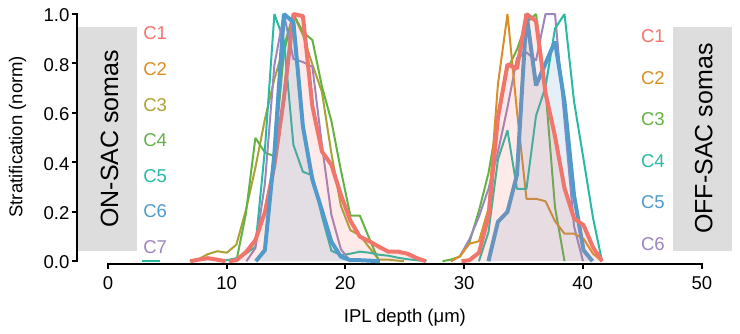

**Figure 6-S2: Stratification profile of the functional clusters detected from motion responses in ON- and OFF-SACs**

Functional clusters whose combination was found to lead to the strongest postsynaptic DS are highlighted in bold and shaded for emphasis.

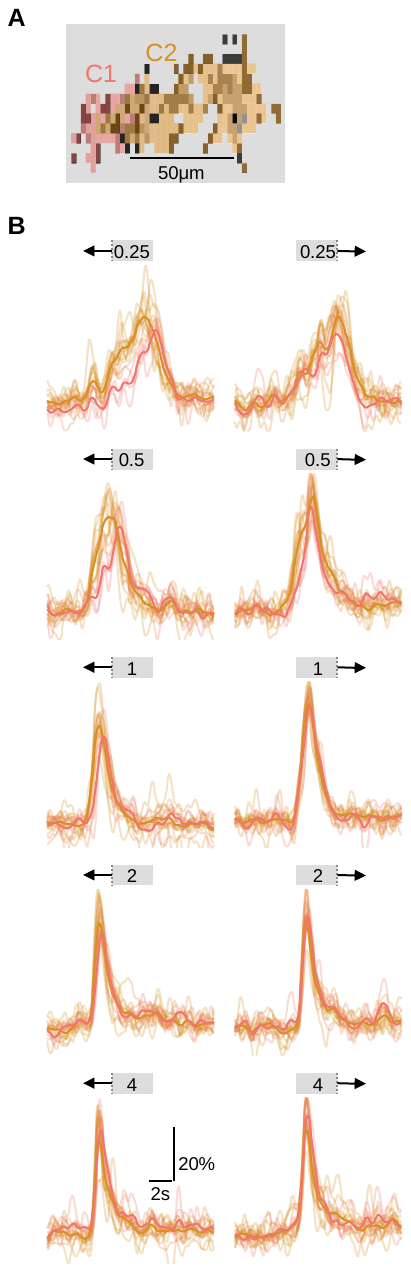

**Figure 6 S3: Comparison of response onset from different functional clusters.**

**A**) A single field of view recorded in ON-SAC dendrites expressing iGluSnFR. Each ROI is color-coded based on its functional cluster identity, displayed in grayscale.

**B**) Changes in fluorescence for different stimuli speeds (from 0.25 mm/s to 4 mm/s) and directions. Color coding indicates functional cluster; bold traces – average responses across all ROIs belonging to the same cluster. The onset of ON-C2 responses was earlier in both directions. This effect is particularly pronounced at slower velocities.

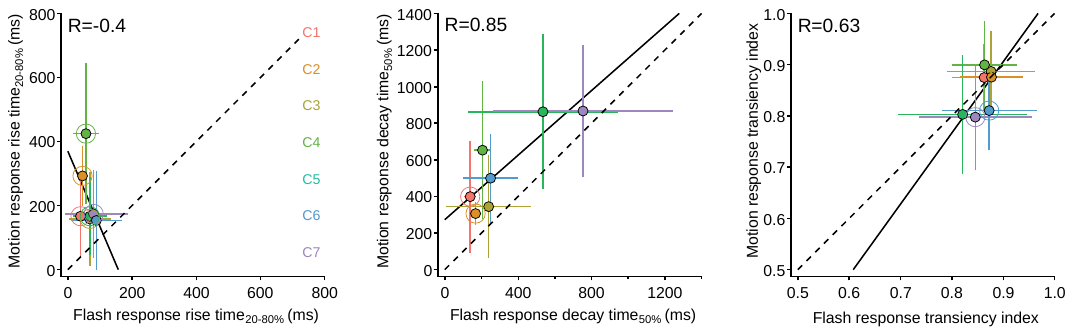

**Figure 7-S1: Comparison of response dynamics to moving bars and stationary flashes.**

Waveform parameters measured from responses recorded in ON-SAC ROIs to moving bars (speed = 0.5 mm/s) plotted as the function of the dynamics in responses to full-field flashes. Black lines represent linear fits. R is the Pearson correlation coefficient. Clusters with significantly different parameters (p<0.05, paired t-test followed by Bonferroni correction for multiple comparisons) are highlighted with larger circles. Error bars represent SD.

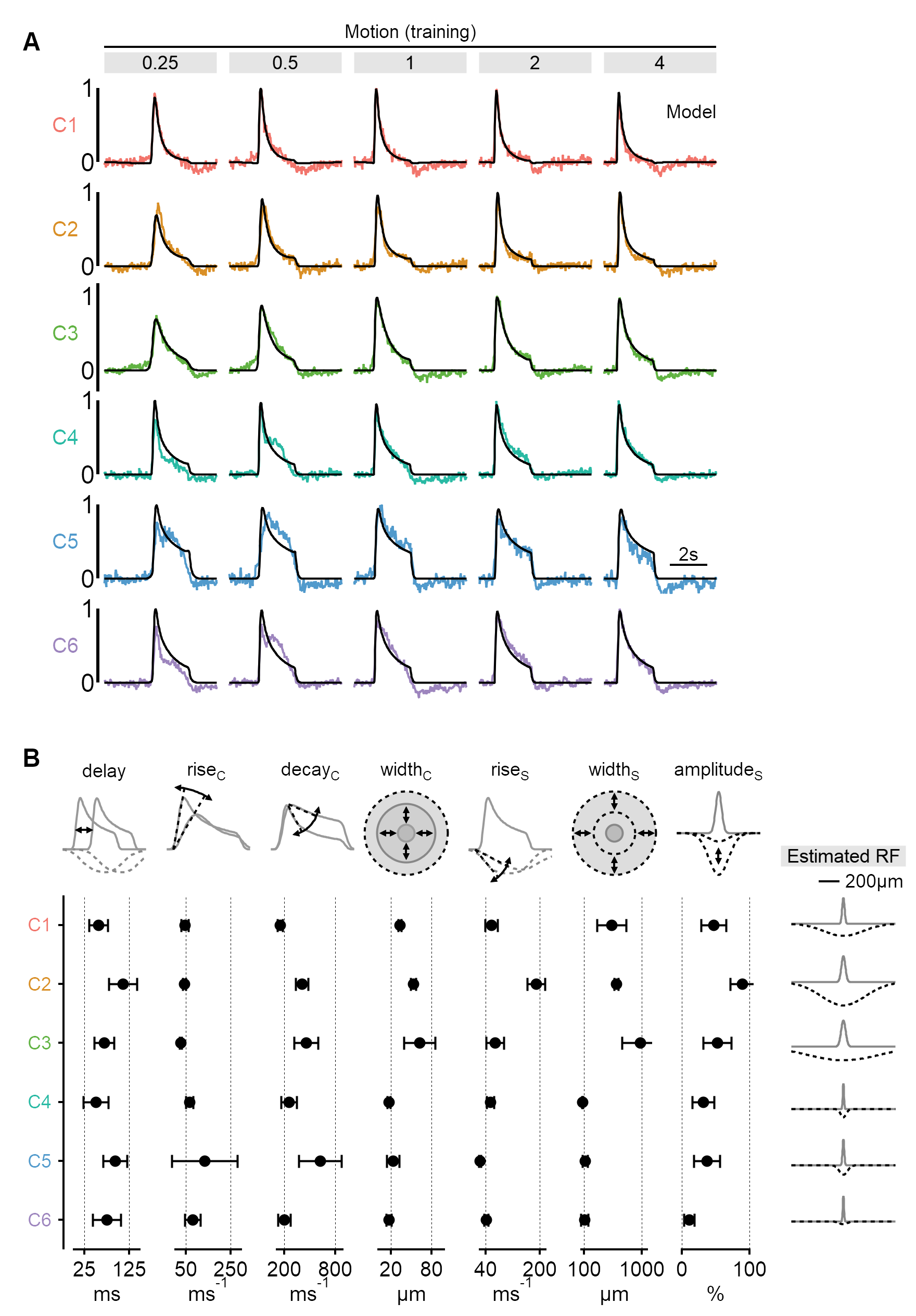

**Figure 8-S1: Estimated RF properties from presynaptic responses to motion in OFF-SACs.**

**A**) As in **Figure 8A**, for functional clusters innervating OFF-SACs.

**B**) As in **Figure 8C**, for the dataset presented in **A**.

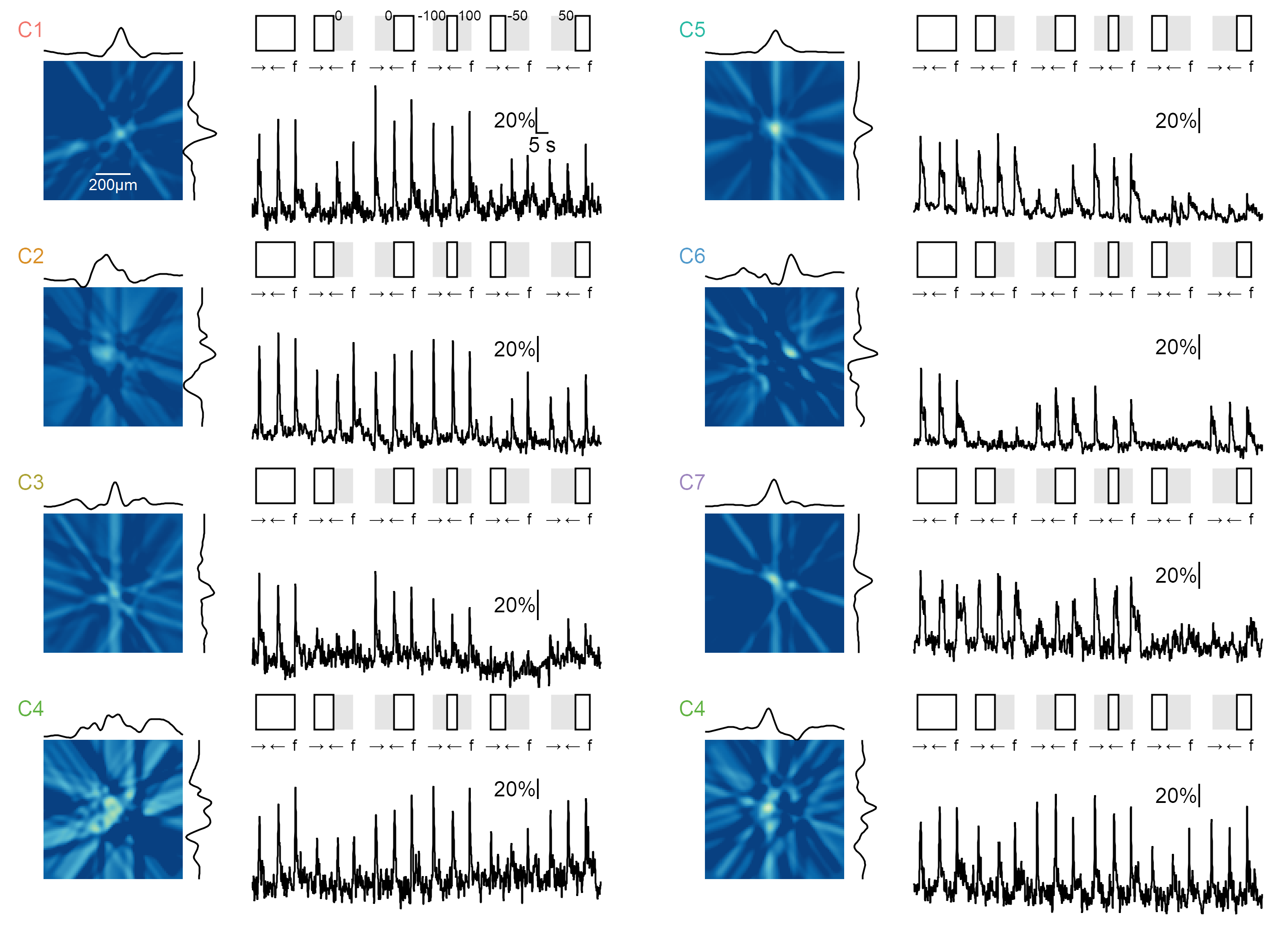

**Figure 7-S3: Example RF shape and responses to moving and static stimuli with full-field and masked stimulation.**

For each ON cluster, the left plot displays a representative RF shape obtained using the filter back projection technique. The black curves represent the x and y RF profiles measured at the center of mass.

On the right, the responses in the same ROI are shown for a series of stimuli consisting of rightward and leftward motion (speed = 0.5 mm/s) and flashes ('f', duration = 2 s) under the following conditions (illustrated schematically at the top): full-field stimulation, masks over the right/left halves of the visual arena, stimuli confined to a region extending 100 µm from the horizontal center of the arena, and right/left masks with edges 50 µm from the horizontal center of the arena. Two examples are shown for the ON-C4 cluster.

The responses to flashes generally align with the spatial extent of the RF as determined using the filtered back projection technique. Note the asymmetry in motion responses near edges.

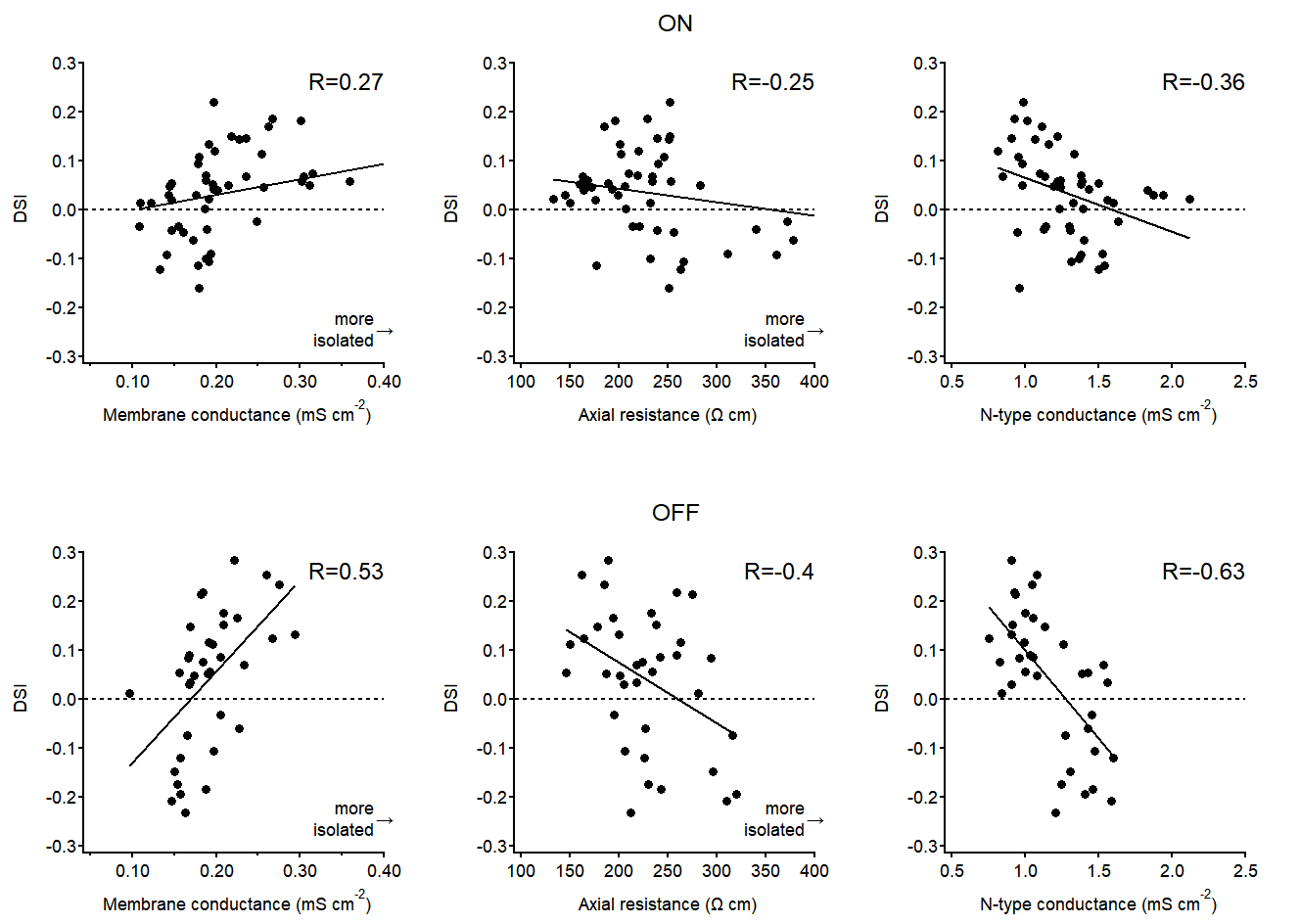

**Figure 9-S1: Correlation between directional tuning and postsynaptic passive and active parameters.**

Directional tuning of each proximal-distal pair shown in **Figure 9** as a function of membrane conductance (reciprocal of membrane resistance, left), axial resistance (middle) and the voltage-gated calcium conductance (right) measured in the optimal model. Lines, linear fits. Higher DS levels were associated with a leakier membrane and lower internal resistance in SACs. These parameters have opposite effects on dendritic isolation.

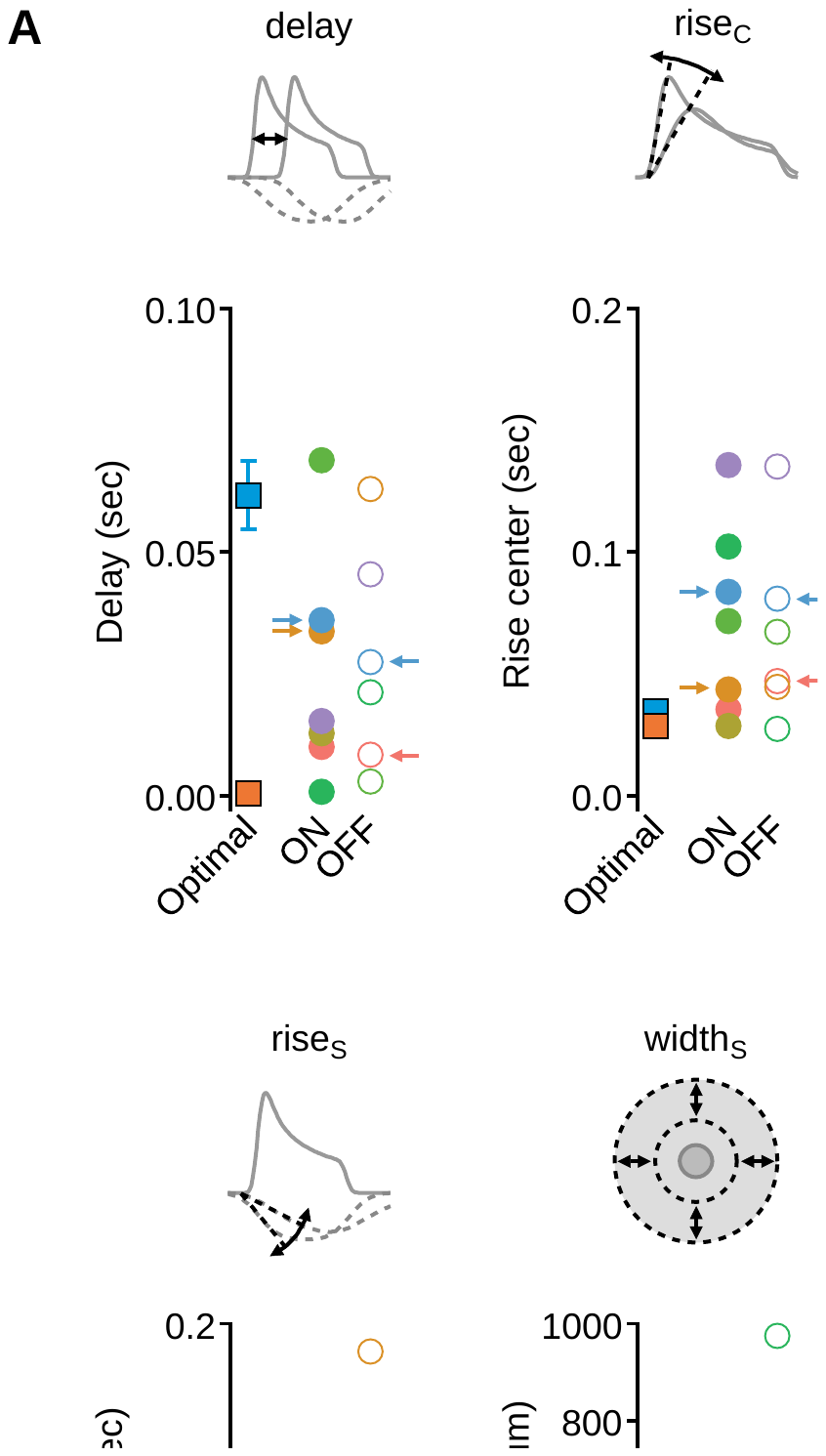

**Figure 11-S1: Comparison of receptive field structure between experimentally determined clusters and the synthetic model mediating optimal directional sensitivity.**

**A**) The mean (±SD) values measured for RF synthetic and experimental components. Data presented as in **Figure 11**.

**B**) Comparison of response waveforms as a function of stimulus velocities produced by optimal synthetic model (top, same data as in **Figure 2**) and experimentally recorded glutamate signals (middle, bottom, same data as in **Figure 10**).
